# Progression of vascular function and blood pressure in a mouse model of Kawasaki disease

**DOI:** 10.1101/2022.08.25.505303

**Authors:** Mia DelVechio, Juliano V Alves, Alina Z Saiyi, Shubhnita Singh, Joseph Galley, Wanessa M. C. Awata, Ariane Bruder-Nascimento, Thiago Bruder-Nascimento

## Abstract

**Background:** Kawasaki disease (KD) is a systemic vasculitis of childhood characterized by vascular damage in the acute stage, which can persist into the late stage. Aneurysms, myocardial infarction, and death are long-standing complications associated with KD. The vascular mechanisms in the cardiovascular risk of KD are not fully studied. Herein, we investigated the vascular function and blood pressure regulation in a murine model of KD.

**Material and methods:** We used the *Candida albicans* water soluble (CAWS) fraction model. Mice were injected with 4 mg CAWS for five consecutive days and separated into 3 groups. Control (Ctrl): water injected for 5 days; CAWS 7 days (C7): CAWS injected for 5 days plus 2 additional days of wait; CAWS 28 days (C28): CAWS injected for 5 days plus 23 additional days of wait. Blood pressure was analyzed via radiotelemetry. In the end, the heart and arteries were harvested for vasculitis characterization and vascular function in a wire myograph. Rat Aortic Vascular Smooth Muscle cells (RASMC) were used to dissect the molecular mechanisms *in vitro*.

**Main findings:** C7 presented elevated inflammatory markers in the coronary area (CA) and abdominal aortae (AA), whereas C28 showed severe vasculitis in CA and AA. No difference was found in blood pressure and heart rate. Vascular dysfunction characterized by higher contractility to norepinephrine (NA) in C7 and C28 was abolished by blocking nitric oxide (NO) production, reactive oxygen species (ROS), and cyclooxygenase (COX)-derived products. RASMC treated with CAWS (10ug/mL) presented an increase in COX2 expression, which was prevented by pre-treating the cells with TAK-242 [Toll like receptor 4 (TLR4) antagonist, 3_x_10^−5^M].

**Conclusion:** Our data indicate that the murine model of KD is associated with vascular dysfunction likely dependent on COX-derived products, oxidant properties, and NO bioavailability. Furthermore, VSMC may present an important role in the genesis of vascular dysfunction and vasculitis via the TLR4 pathway. Finally, the CAWS model seems not to be appropriate to study KD-associated shock. More studies are necessary to understand whether vascular dysfunction and COXs are triggers for vasculitis.

## Introduction

Kawasaki disease (KD) is a severe multisystem inflammatory syndrome characterized by vasculitis^1^, which affects infants and young children^2,3^. KD is the leading cause of acquired cardiovascular disease among children in North America, Europe, and Japan^1,3,4^. It affects approximately 9 to 20 children per 100,000^4^. If not diagnosed or treated properly, KD can result in permanent cardiovascular sequelae that extend into adulthood^5-7^. Coronary arteritis followed by coronary artery aneurysms is a major cardiovascular risk of KD and it can occur in up to 30% of untreated children^8,9^. High-dose intravenous immunoglobulin (IVIG) seems to be the most efficient treatment to reduce cardiovascular injury in kids with KD, however resistant cases to IVIG are associated with a higher risk for cardiovascular complications including aneurysms and Kawasaki disease shock syndrome (KDSS)^10-16^, which is characterized by a 20% decrease or more in systolic blood pressure compared to healthy individuals at the same age^10^.

A wide variety of innate and adaptive immune cells, including monocytes, macrophages, neutrophils, and T-lymphocytes, infiltrate the coronary artery wall in KD, which is linked to the condition ^3,17-22^. These cells produce a large quantity of inflammatory cytokines such as Interleukin 1β (IL1β), tumor necrosis factor a (TNFα), and Interleukin 17 (IL17) that can directly disturb the vascular homeostasis^23-25^ and lead to vascular dysfunction as described elsewhere^23-26^. Vascular dysfunction is an important event in the genesis of several cardiovascular risks including hypertension and atherosclerosis^27-29^. Nitric oxide (NO), reactive oxygen species (ROS), and cyclooxygenase (COX)-derived products (prostaglandins), which are endothelium derived factors, are master regulators of vascular function via regulating changes in vascular smooth muscle cells (VSMC) contractile characteristics and arterial stiffness. ^23,30-33^.

Young people and adults with a history of KD have previously been reported to have vascular dysfunction, which increases the risk of coronary vascular disturbances years after KD recovery. ^6,34-37^. In mouse model of KD, early exposition to KD inducer facilitates the atherosclerosis appearance later in life, indicating that vascular dysfunction (a key event on the genesis of atherosclerosis) is present in KD^38^. However, few other studies reported that vascular dysfunction in KD does not persist into adulthood^9,39^. The molecular underpinnings of vascular dysfunction are completely unknown or speculative, despite the fact that it has already been observed in humans. Therefore, mouse models of vasculitis are useful tools, especially for analyzing the molecular and cellular mechanisms underlying KD-related vascular dysfunction and for assessing vascular health both during acute and convalescent phases.

In this study, we aimed to determine if the acute and convalescent phases of KD in a mouse model are related with vascular dysfunction as well as to dissect the molecular processes behind any change in vascular function. We also tried to characterize if a murine model of KD could be a good approach to study KDSS. Thus, we tested the hypothesis that the *Candida albicans* water soluble (CAWS) model of KD is associated with vascular dysfunction and changes in blood pressure regulation.

### Animals

Six- to twelve-week-old male C57BL6/J wild type were used. All mice were fed with standard mouse chow and tap water was provided ad libitum. Mice were housed in an American Association of Laboratory Animal Care– approved animal care facility in Rangos Research Building at Children’s Hospital of Pittsburgh of University of Pittsburgh. Institutional Animal Care and Use Committee approved all protocols (IACUC protocols # 19065333 and 22061179). All experiments were performed in accordance with Guide Laboratory Animals for The Care and Use of Laboratory Animals.

### Preparation of Candida albicans Water Soluble (CAWS) fraction

The CAWS complex was prepared as described elsewhere^21,22,40,41^. Briefly, *C. albicans* ATCC 18804^®^ was cultured in C-limiting medium^42^ for 48h. An equal volume of ethanol was added and allowed to stand overnight. The mixture was centrifuged, and the pellet was dissolved in water and centrifuged again. An equal volume of ethanol was added to the soluble fraction and the mixture was left to stand overnight. The mixture was centrifuged, and the precipitate was dried with acetone. The CAWS complex was dissolved in water and autoclaved. Aliquots were maintained in -20ºC freezer.

### CAWS-Induced Vasculitis Model of Mice

Mice were injected with 4 mg CAWS (i.p.) for five consecutive days. Animals were separated into 3 groups. Control: water injected for 5 days; CAWS 7 days: CAWS injected for 5 days plus 2 additional days of wait; CAWS 28 days: CAWS injected for 5 days plus 23 additional days of wait^21^ (Fig. 1A). At the end of these periods, mice were killed by CO_2_ saturation followed by decapitation.

**Figure 1.**
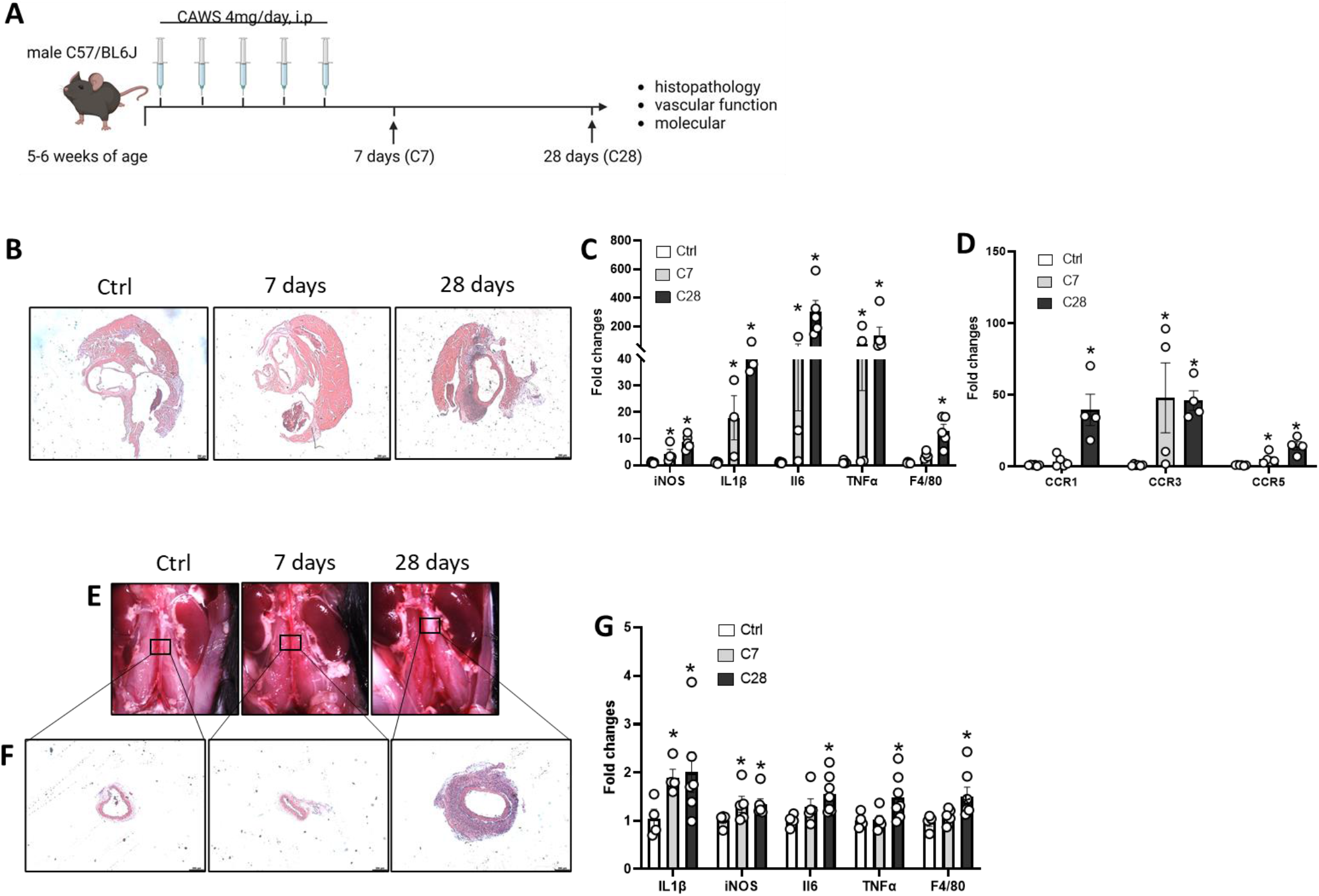
CAWS induces a severe vasculitis in coronary area and abdominal aorta. Schematic of Kawasaki disease induction (A). Vasculitis levels in aortic root and coronary areas (B), cardiac inflammatory (C) and cardiac chemokines receptors (D) gene expression. Vasculitis in abdominal aortae (E-F) and aortic inflammatory gene expression from control (Ctrl): water injected for 5 days; CAWS 7 days (C7): CAWS injected for 5 days plus 2 additional days of wait; CAWS 28 days (C28): CAWS injected for 5 days plus 23 additional days of wait. N=5-7. Data are presented as Mean ± SEM. *P<0.05 vs Ctrl. Images represent a single experiment from 5 repetitions. Ruler length: 200uM

### Characterization of vasculitis

Mice were euthanized for heart and aortae harvest at day 7- or 28-days post CAWS and perfused with cold phosphate-buffered saline (PBS). Tissues were collected and placed in 4% paraformaldehyde (PFA) solution for histology analysis or in cold Krebs Henseleit Solution for vascular function studies. Abdominal aortas were photographed in the animal and scored for vasculitis. After 12h in PFA, tissues were placed in 70% ethanol until the day of preparing the samples for histology. Hearts and aortae were embedded in paraffin, then samples were sectioned and stained with hematoxylin and eosin (H&E) to analyze the vasculitis and damage caused by KD. The samples were blinding scored for vasculitis appearance.

### *In vivo* blood pressure measurement

When compared to healthy people of the same age, KDSS patients’ systolic blood pressure often drops by more than 20%. ^10,15,16^. Although it is still uncommon, 1-7% of KD cases have it^10^. Herein, we sought to characterize whether CAWS-induced KD might change the blood pressure. Mice with 6-7 weeks of age were control or HFD mice were instrumented with telemetry transmitters to record arterial pressure and heart rate (HD-X10, Data Sciences International, St Paul, MN). Transmitters were implanted as described previously^43,44^. After 7 days of recovery from surgery, necessary for the mice to gain their initial body weight, data were recorded for 5 days as baseline. Then, KD was induced as described before, and blood pressure and heart rate values were obtained at 10-min intervals for the duration of 7 days post-first injection of CAWS.

### Indices of autonomic function

To analyze whether changes in autonomic response interfere with any changes in blood pressure, indices of the autonomic function were obtained on the last day of the recording baseline period and after 7 days post-first injection of CAWS. A classic pharmacological method consisting in a single intraperitoneal injection of the ganglionic blocker mecamylamine (5 mg/kg) or of the β-adrenergic receptor blocker propranolol (6 mg/kg) was used. Injections were conducted more than 2 h apart in a random order. Changes in blood pressure or heart rate response to pharmacological compounds within 60 min post-injection were reported. Changes in mean arterial pressure (MAP) or heart rate (HR) were expressed as a percent of the baseline MAP or heart rate^43,44^.

### Vascular function

Since abdominal aortae are prone to develop severe vasculitis^1,19,45^ in a mouse model of KD, we used sections of these arteries to study the vascular function. Briefly, abdominal aortae were dissected from connective tissues, separated into rings (1.5-2mm), and mounted in a wire myograph (Danysh MyoTechnology) for isometric tension recordings with PowerLab software (AD Instruments). Rings were placed in tissue baths containing warmed (37 °C), aerated (95% O_2_, 5% CO_2_) Krebs Henseleit Solution: (in mM: 130 NaCl, 4.7 KCl, 1.17 MgSO_4_, 0.03 EDTA, 1.6 CaCl_2_, 14.9 NaHCO_3_, 1.18 KH_2_PO_4_, and 5.5 glucose). After 30 min of stabilization, KCL (120 mM) was used to test arterial viability. Concentration-response curves (CRC) for norepinephrine (NA) were performed in presence of COX1-2 inhibitor (Indomethacin, 10^−5^M), COX2 inhibitor (NS398, 10^−5^M), thromboxane A_2_ (TXA_2_) receptor antagonist (NT-126, 10^−5^M), Superoxide dismutase (SOD) mimetic (Tempol, 10^−4^M), and NOS inhibitor (L-NAME, 10^−4^M)^32,44,46^. CRC to U46619 (TXA_2_ analogue) was used to analyze whether any difference in NA response was an exclusive adrenergic dependent response.

### Western Blot

RASMC samples were directly homogenized using 2x Laemmli Sample Buffer and supplemented with 2-Mercaptoethanol (β-mercaptoethanol) (BioRad Hercules, California – USA). Aortic protein was extracted using radioimmunoprecipitation assay buffer (RIPA) buffer (30mM HEPES, pH 7.4,150mM NaCl, 1% Nonidet P-40,0.5% sodium deoxycholate, 0.1% sodium dodecyl sulfate, 5mM EDTA, 1mM NaV04, 50mM NaF, 1mM PMSF, 10% pepstatin A, 10 μg/ml leupepsin, and 10 μg/ml aprotinin). Proteins were separated by electrophoresis on a polyacrylamide gradient gel (BioRad Hercules, California – USA), and transferred to Immobilon-P poly (vinylidene fluoride) membranes. Non-specific binding sites were blocked with 5% skim milk or 1% bovine serum albumin (BSA) in tris-buffered saline solution with tween for 1h at 24 ºC. Membranes were then incubated with specific antibodies overnight at 4 ºC as described in table 1. After incubation with secondary antibodies, the enhanced chemiluminescence luminol reagent (SuperSignal™ West Femto Maximum Sensitivity Substrate, Thermo Fisher Waltham, MA, USA) was used for antibody detection.

**Table 1.** Antibodies list

| Antibody | Manufacturer | Catalog number | Concentration | Secondary |
| --- | --- | --- | --- | --- |
| COX1 | Cell Signaling | 4841 | 1:1000 | rabbit |
| COX2 | BD Biosciences | 610203 | 1:500 | mouse |
| Nox1 | Sigma | SAB4200097 | 1:1000 | rabbit |
| Nox2 | Abcam | Ab129068 | 1:1000 | rabbit |
| Nox4 | Abcam | Ab133303 | 1:1000 | rabbit |
| Ser<sup>1177</sup> eNOS | BD Biosciences | 612393 | 1:1000 | mouse |
| eNOS | BD Biosciences | 610296 | 1:1000 | mouse |
| iNOS | Cell Signaling | 13120 | 1:500 | rabbit |
| βactin | Sigma | A3854 | 1:20000 | HRP conjugated |
| GAPDH | Cell Signaling | 3683 | 1:1000 | HRP conjugated |

HRP: horseradish peroxidase

### RT-PCR

mRNA from aortae, hearts, and VSMC was extracted using RNeasy Mini Kit (Quiagen, Germantown, MD – USA). Complementary DNA (cDNA) was generated by reverse transcription polymerase chain reaction (RT-PCR) with SuperScript III (Thermo Fisher Waltham, MA USA). Reverse transcription was performed at 58 ºC for 50 min; the enzyme was heat inactivated at 85 ºC for 5 min, and real-time quantitative RT-PCR was performed with the PowerTrack™ SYBR Green Master Mix (Thermo Fisher, Waltham, MA USA). Sequences of genes as listed in table 2. Experiments were performed in a QuantStudio™ 5 Real-Time PCR System, 384-well (Thermo Fisher, Waltham, MA USA). Data were quantified by 2ΔΔ Ct and are presented by fold changes indicative of either upregulation or downregulation.

**Table 2.**
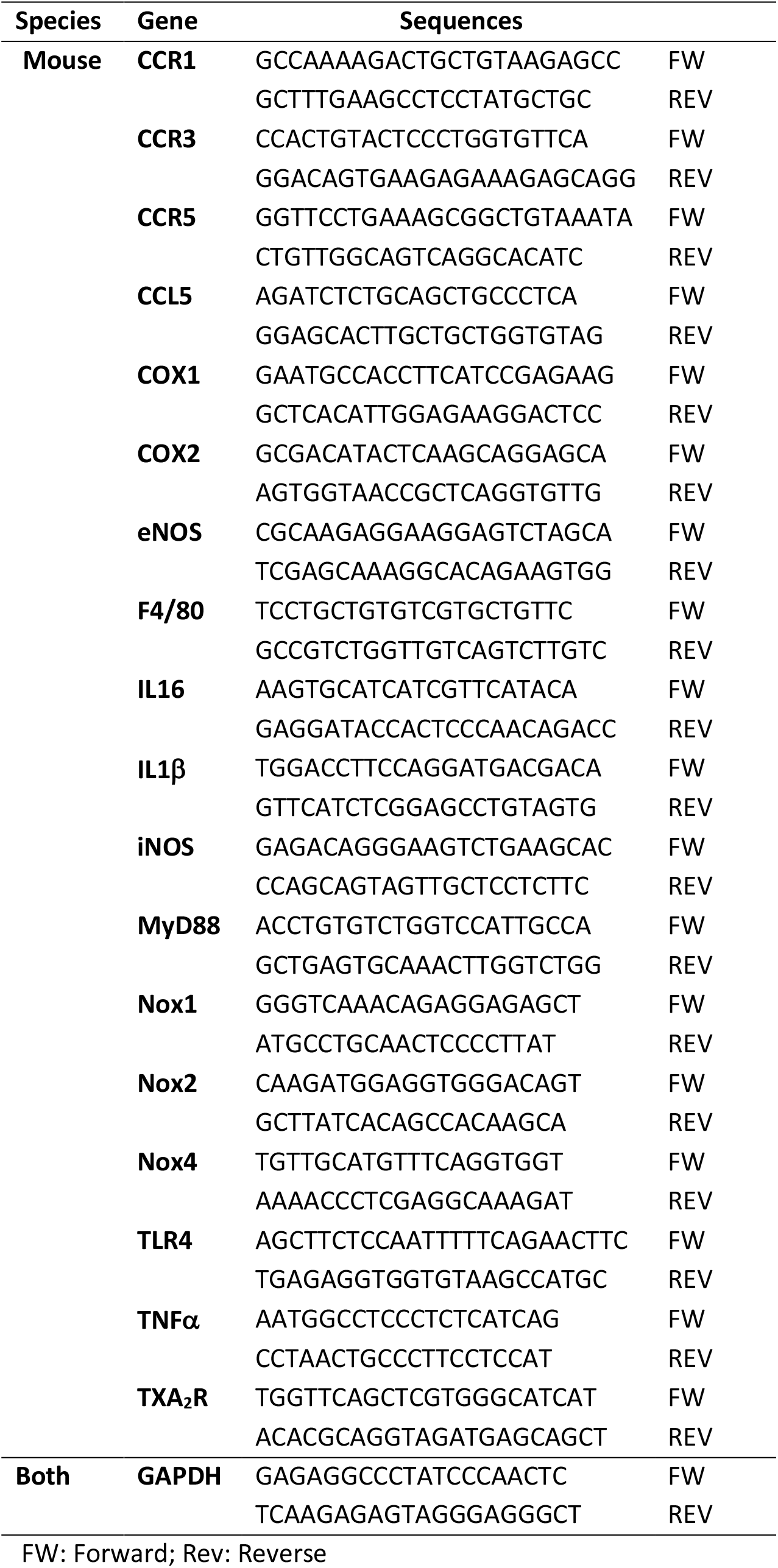
List of primers

### Culture of Vascular Smooth Muscle Cells

As described by our previous studies^26,33^, Rat Aortic Smooth Muscle cells (RASMC, Lonza, Walkersville, MD, USA) were maintained in Dulbecco’s Modified Eagle Medium (DMEM) (Gibco from Thermo Fisher Scientific, Waltham, MA, USA) containing 10% fetal bovine serum (FBS, HyClone Logan UT, USA), 100 U/ml penicillin and 100 μg/ml streptomycin (Gibco from Thermo Fisher Scientific, Waltham, MA, USA).

### Protocol of VSMC treatment

The complete medium was replaced by DMEM containing 0.5% FBS before the treatments. Cells were treated with CAWS (10ug/mL) in different times (0-24h). To understand whether TLR4 can recognize CAWS to generate vascular changes, we pre-incubated TLR4 Inhibitor (TAK-242, 3_x_10^−5^M, Sigma, St. Louis, MO, USA) 30 minutes before CAWS treatment. Antibodies used are described in table 1.

### Statistic

For comparisons of multiple groups, one-way or two-way analysis of variance (ANOVA), followed by the Tukey post-test was used. Differences between two groups were determined using Student’s t test. The vascular function findings are expressed as absolute numbers (mN). The concentration-response curves were fitted by nonlinear regression analysis. pD2 (defined as the negative logarithm of the EC50 values) and maximal response (Emax) were determined. Analyses were performed using Prism 9.0 software (GraphPad). A difference was considered statistically significant when P ≤ 0.05.

## Results

### CAWS induces a severe vasculitis

We first analyzed the levels of vasculitis in our mouse model of KD 7- and 28-days post first injection of CAWS (Fig. 1A). At 7 days (C7 group), we did not observe any signal of vasculitis by histology in coronary area (Fig. 1B) or abdominal aortae (Fig. 1E-F), however we did see an increase in several inflammatory genes in heart and abdominal aortae including iNOS, F4/80 (macrophage marker), IL-1β, IL-6, and TNFα, as well as increase in CCR3 and CCR5 (Fig. 1B-D). No difference was observed in CCR1 in C7 group. At 28 days (C28 group), a severe vasculitis in coronary area was found (Fig. 1B) and abdominal aortae (Fig. 1E-F) with a striking elevation in inflammatory genes (iNOS, F4/80, IL-1β, IL-6, and TNFα).

### CAWS treatment does not affect blood pressure and autonomic responses

Although KDSS is considered a rare characteristic in acute phase of KD (REF), we analyzed the MAP and heart rate via telemetry in acute phase of KD to analyze whether our mouse model of KD is appropriate to study KDSS. We observed a mild reduction in systolic, diastolic, and MAP, (not significant) with no changes in HR during the CAWS treatment (Figure 2B-E). Furthermore, we analyzed the involvement of sympathetic tone before and after CAWS treatment by injecting the mice with a β-blocker or a ganglionic blocker as described elsewhere^43^. We found that the drop in HR and MAP caused by propranolol and mecamylamine respectively persisted similarly before and after KD induction (Fig. 2F-G). These findings indicate that this mouse model of KD is not appropriate to study KDSS, and it does not affect the sympathetic tone.

**Figure 2.**
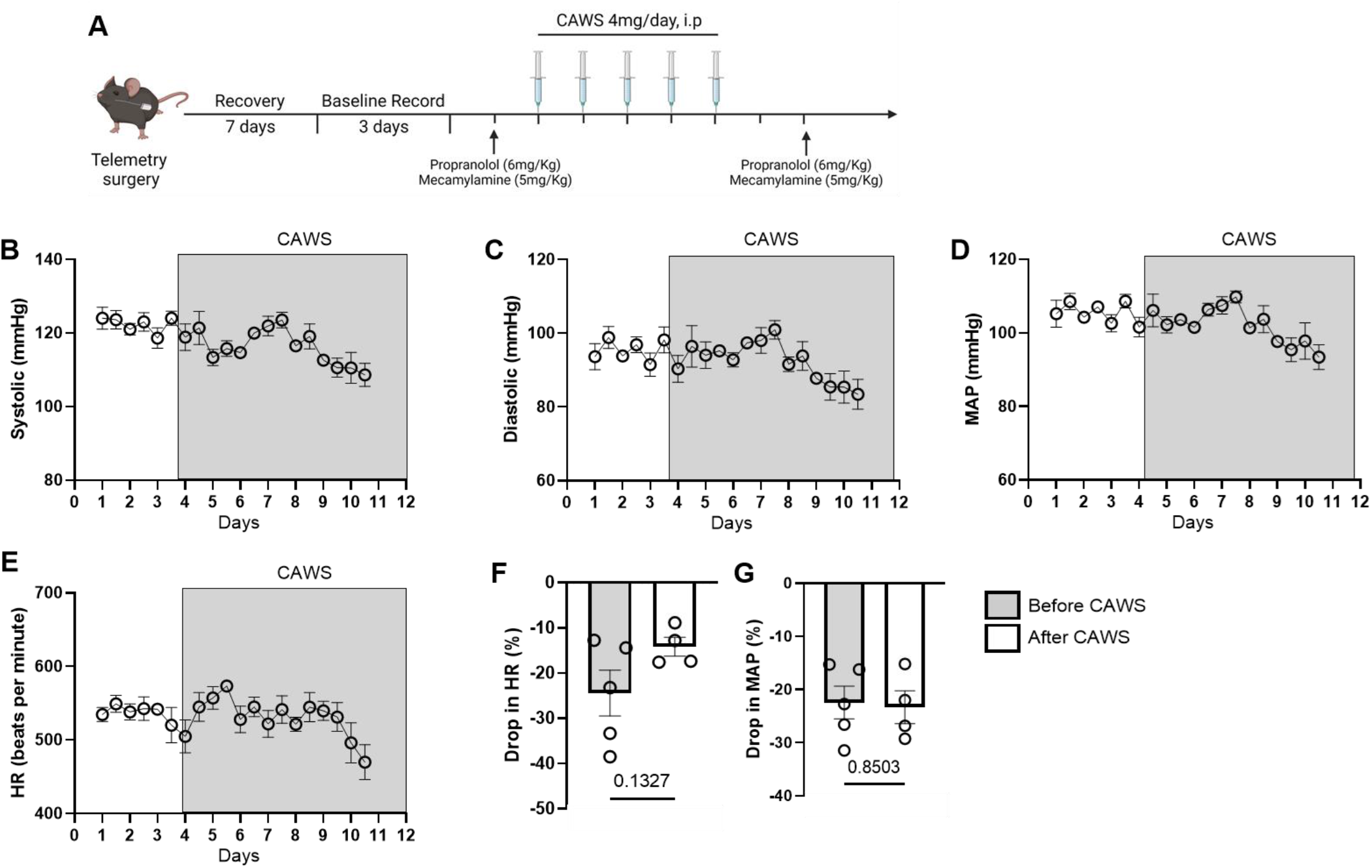
CAWS-induced vasculitis does not change blood pressure. Schematic of blood pressure measurement and Kawasaki disease induction (A). Systolic blood pressure (B), diastolic blood pressure (C), mean arterial pressure (MAP) (D) and heart rate (HR) (E) measured by radiotelemetry (3 days as baseline and 7 days as Kawasaki disease). Effects of propranolol (6mg/kg) on HR (F) and effects of mecamylamine (5mg/Kg) on MAP (G) of mice before and after treatment with CAWS. CAWS was injected for 5 days plus 2 additional days of wait. N=4-5. Data are presented as Mean ± SEM. *P<0.05 vs Ctrl. Difference in number of experiments in propranolol and mecamylamine effects before and after CAWS treatment occurred because a mouse died after propranolol injection post-CAWS treatment.

### KD is associated with changes in vascular contractility via regulating COXs-derived products

We analyze via wire myograph the vascular contractility in abdominal aortae (a sensitive artery to the effects of CAWS) from our KD model at 7 and 28 post first CAWS injection. Strikingly, we observed that arteries from C7 and G28 presented an increased maximal response to NA compared to Ctrl group (Fig. 3 and Table 3), with no difference sensitive (Table 3). No difference was observed between C7 and G28 (Fig. 3 and Table 3).

**Table 3.** Maximal response and pD2

| Groups | Parameters |  |
| --- | --- | --- |
|  | Maximal response (Emax) | pD2 |
| Ctrl | 3.045 ± 0.46 | 6.002 ± 0.18 |
| C7 | 5.098 ± 0.34* | 6.610 ± 0.42 |
| C28 | 5.386 ± 0.71* | 6.561 ± 0.30 |

Control (Ctrl): water injected for 5 days; CAWS 7 days (C7): CAWS injected for 5 days plus 2 additional days of wait; CAWS 28 days (C28): CAWS injected for 5 days plus 23 additional days of wait. pD2: negative logarithm of the EC50 values. N=5-7. Data are presented as Mean ± SEM. \*P<0.05 vs Ctrl.

**Figure 3.**
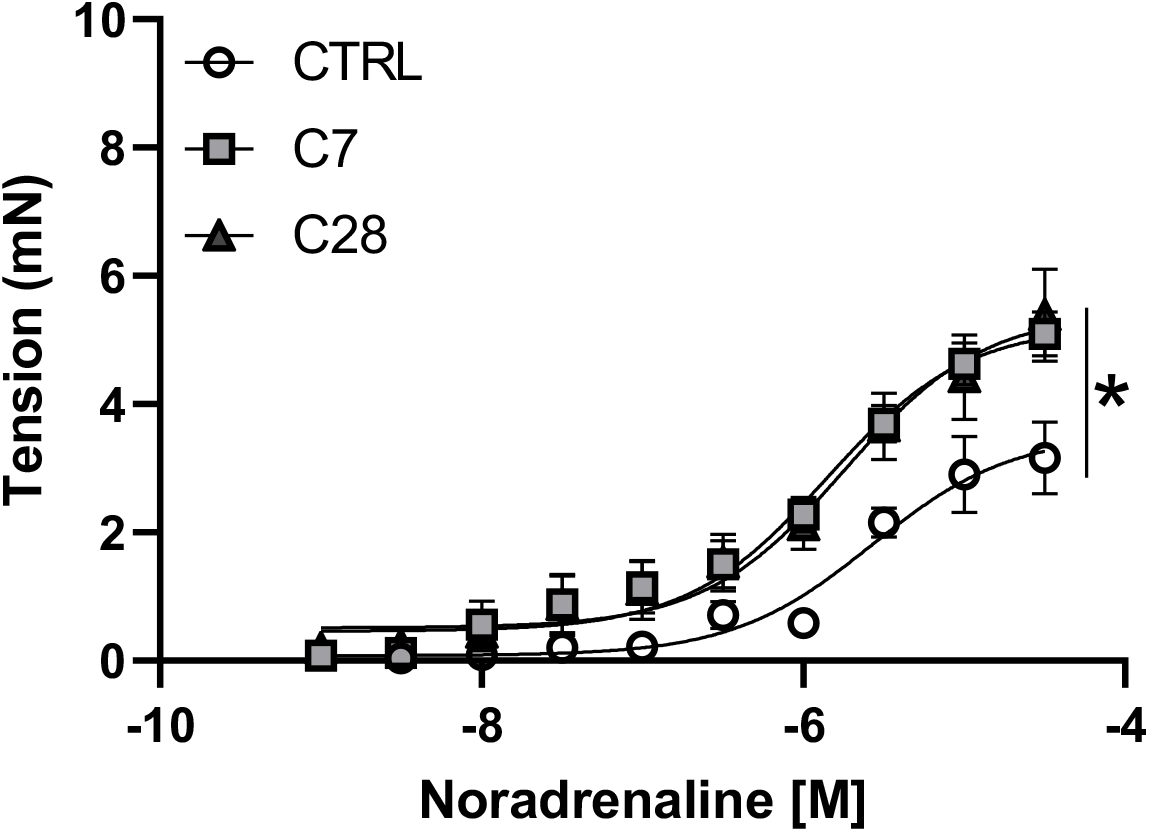
Murine model of Kawasaki disease is associated with vascular dysfunction. Concentration-Response Curves (CRC) to Noradrenaline in abdominal aorta from control (Ctrl): water injected for 5 days; CAWS 7 days (C7): CAWS injected for 5 days plus 2 additional days of wait; CAWS 28 days (C28): CAWS injected for 5 days plus 23 additional days of wait. N=5-7. Data are presented as Mean ± SEM. *P<0.05 vs Ctrl.

To dissect the mechanisms whereby KD is associated with vascular dysfunction, we performed a set of experiments in the presence of different inhibitors. We first blocked COX1 and 2 by pre-treating the arteries with indomethacin and found that COXs inhibition abrogated the vascular differences (Fig. 4A). Next, we blocked only COX2 using a specific inhibitor (NS398) and found that COX2 inhibition restored the vascular contractility like the Ctrl group (Fig. 4B). TXA_2_ is a potent vasoconstrictor derived from COXs^47^. We tested whether TXA_2_ could be leading to the vascular hypercontractility in the KD mouse model by antagonizing its receptor, we interestingly observed that blockage of the TXA_2_ receptor abolished the difference between Ctrl and KD groups (Fig. 4C). We also found that C7 and C28 presented elevated COX2 gene and protein expression and TXA_2_R gene expression in aortae, while COX1 was elevated in C28 only at protein levels (Fig. 4D-F).

**Figure 4.**
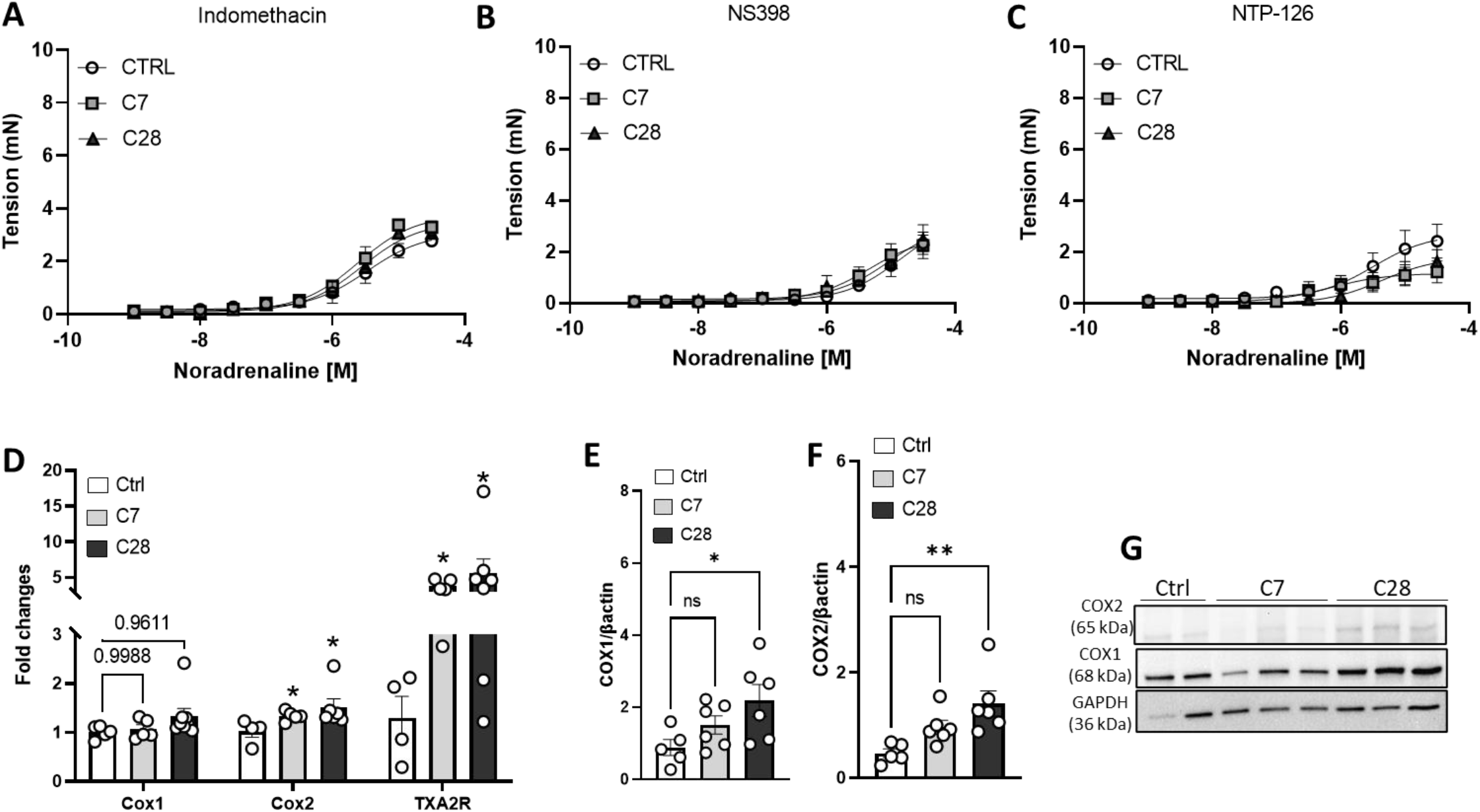
Vascular dysfunction is murine model of Kawasaki disease is dependent on COXs-derived products. Concentration-Response Curves (CRC) to Noradrenaline in abdominal aorta in presence of indomethacin (A), NS398 (B), and NTP-126 (C). Aortic COX1, COX2, and TXA_2_R gene expression (D). Aortic COX1 (E) and COX2 (F) protein expression. Experiments were performed in abdominal aortae from control (Ctrl): water injected for 5 days; CAWS 7 days (C7): CAWS injected for 5 days plus 2 additional days of wait; CAWS 28 days (C28): CAWS injected for 5 days plus 23 additional days of wait. N=5-7. Data are presented as Mean ± SEM. *P<0.05 vs Ctrl.

### NO and ROS contribute to the vascular dysfunction in KD

We studied whether NO and ROS are involved in vascular dysfunction in KD by blocking NOS with L-NAME and Tempol, respectively. L-NAME augmented the vascular response to NA at the same levels in all groups abrogating the difference in maximal response (Fig. 5A). Furthermore, pre-incubation with antioxidant agent restored the vascular contractility in C7 and C28 groups (Fig. 5E). Finally, no difference was observed for eNOS expression at protein and gene levels (Fig. 5B). Increases in Nox1 protein expression, but not in gene, was found in aortae only in C28 group, differently Nox2 was elevated in C7 and C28 in gene and protein expression. No difference was observed for Nox4 (Fig. 5G-I).

**Figure 5.**
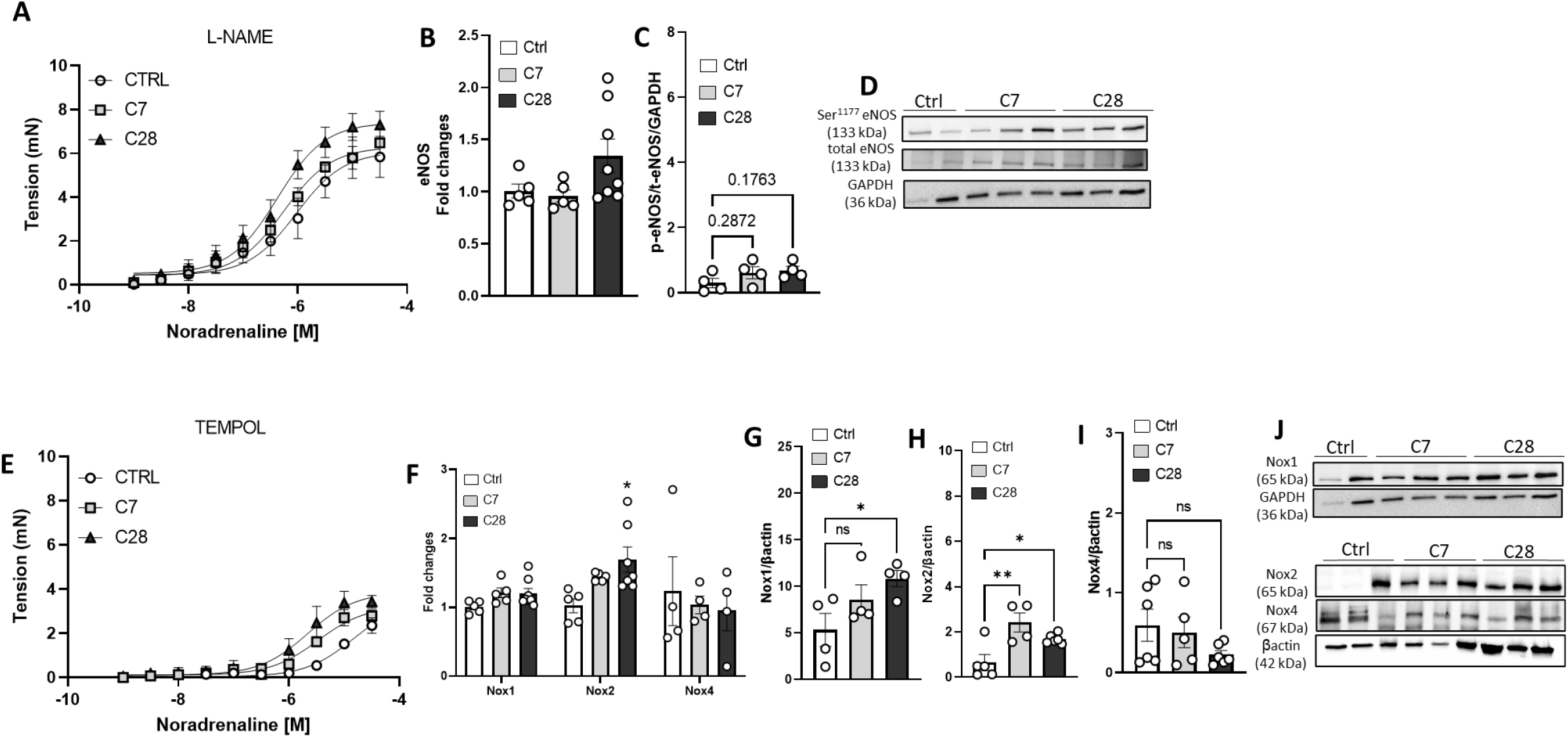
Vascular dysfunction in murine model of Kawasaki disease is dependent on NO and ROS bioavailability. Concentration-Response Curves (CRC) to Noradrenaline in abdominal aorta in presence of L-NAME (A), aortic eNOS gene expression (B), aortic eNOS phosphorylation at Ser^1177^ (C), and aortic iNOS protein expression (D). CRC to Noradrenaline in abdominal aorta in presence of Tempol (E), aortic Nox1, 2, and 4 gene expression (F), and aortic Nox1 (G and J), Nox2 (H and J), and Nox4 (I and J) protein expression. Experiments were performed in abdominal aortae from control (Ctrl): water injected for 5 days; CAWS 7 days (C7): CAWS injected for 5 days plus 2 additional days of wait; CAWS 28 days (C28): CAWS injected for 5 days plus 23 additional days of wait. N=4-7. Data are presented as Mean ± SEM. *P<0.05 vs Ctrl.

### CAWS evokes COXs expression via TLR4 in VSMC

Firstly, we performed a time course with CAWS to analyze whether it can change Nox1 and 4 and COXs expressions in VSMC, we did not analyze Nox2 in RASMC because its expression is very low^33^. We observed no differences in Nox1 and 4 enzymes expression (Fig. 6A-B), but a significant increase in COX1 and 2 at 24h of treatment was found (Fig. 7A-B). Then, we defined 24h stimulus as our main time-point to study how CAWS changes COXs expression in VSMC. TLR4 belongs to the pattern recognition receptor (PRR) family, and it can recognize a pathogen and trigger a very specific cellular response^48^. We observed that MyD88, downstream molecule for TLR4 activation^48,49^, was upregulated in aortae from CAWS treated mice (Fig. 7C). No difference was observed for TLR4 gene expression (Fig. 7C). Thus, we treated VSMC with CAWS with or without TLR4 inhibitor (TAK-242) to analyze whether CAWS can increase COXs via TLR4 in the vasculature. We remarkably found that TLR4 antagonist blocked CAWS-induced COX2 expression in VSMC, although TAK-242 did not significantly block COX1 expression, there was a strong trend (P=0.08) (Fig. 7E-G).

**Figure 6.**
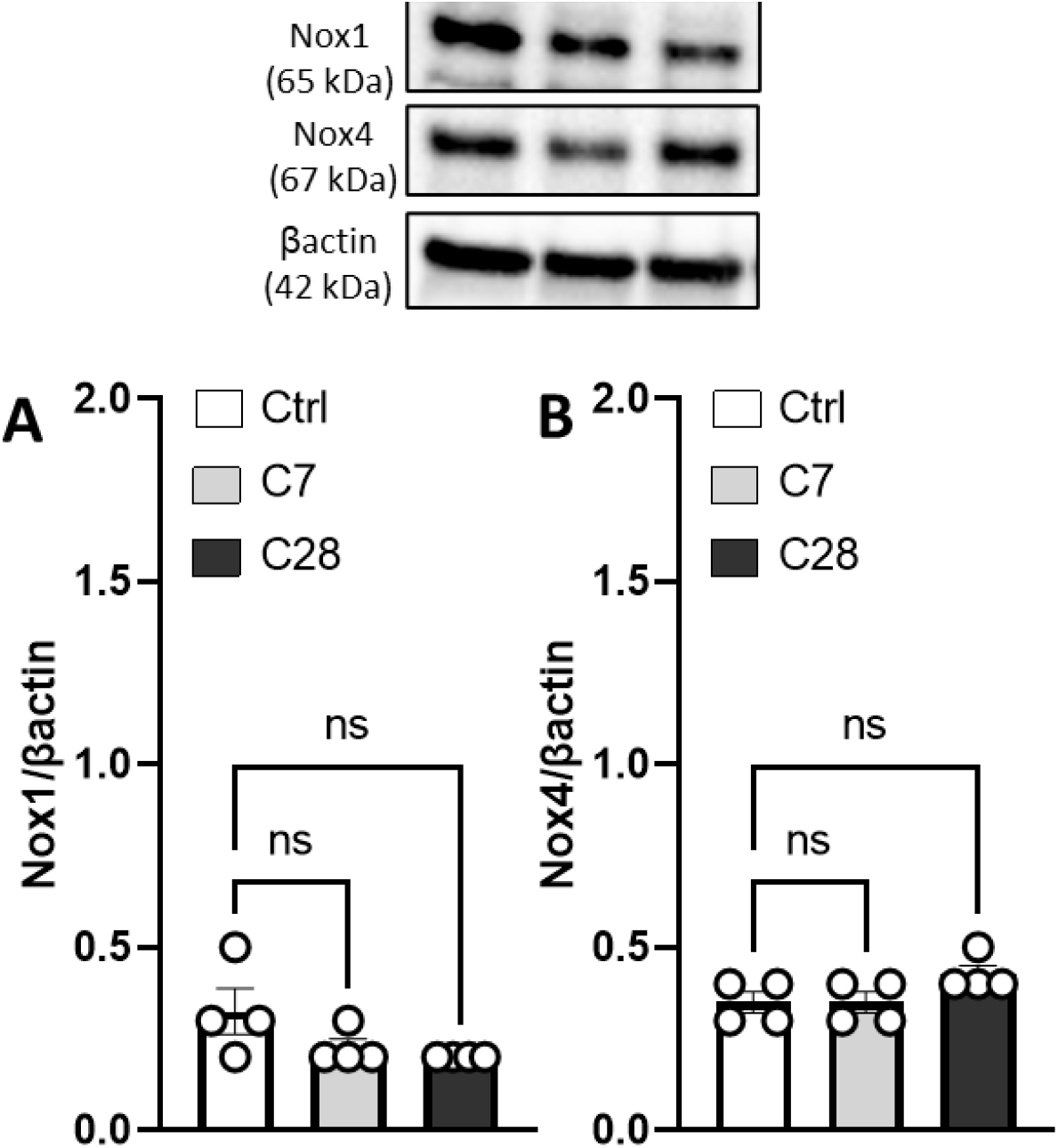
CAWS treatment in vitro does not affect Noxs expression in vascular smooth muscle cells. Effects of CAWS (10ug/mL, 0-24h) on Nox1 (A), and Nox4 (B) protein expression in Rat Aortic Smooth Muscle Cells (RASMC). N=4. Data are presented as Mean ± SEM.

**Figure 7.**
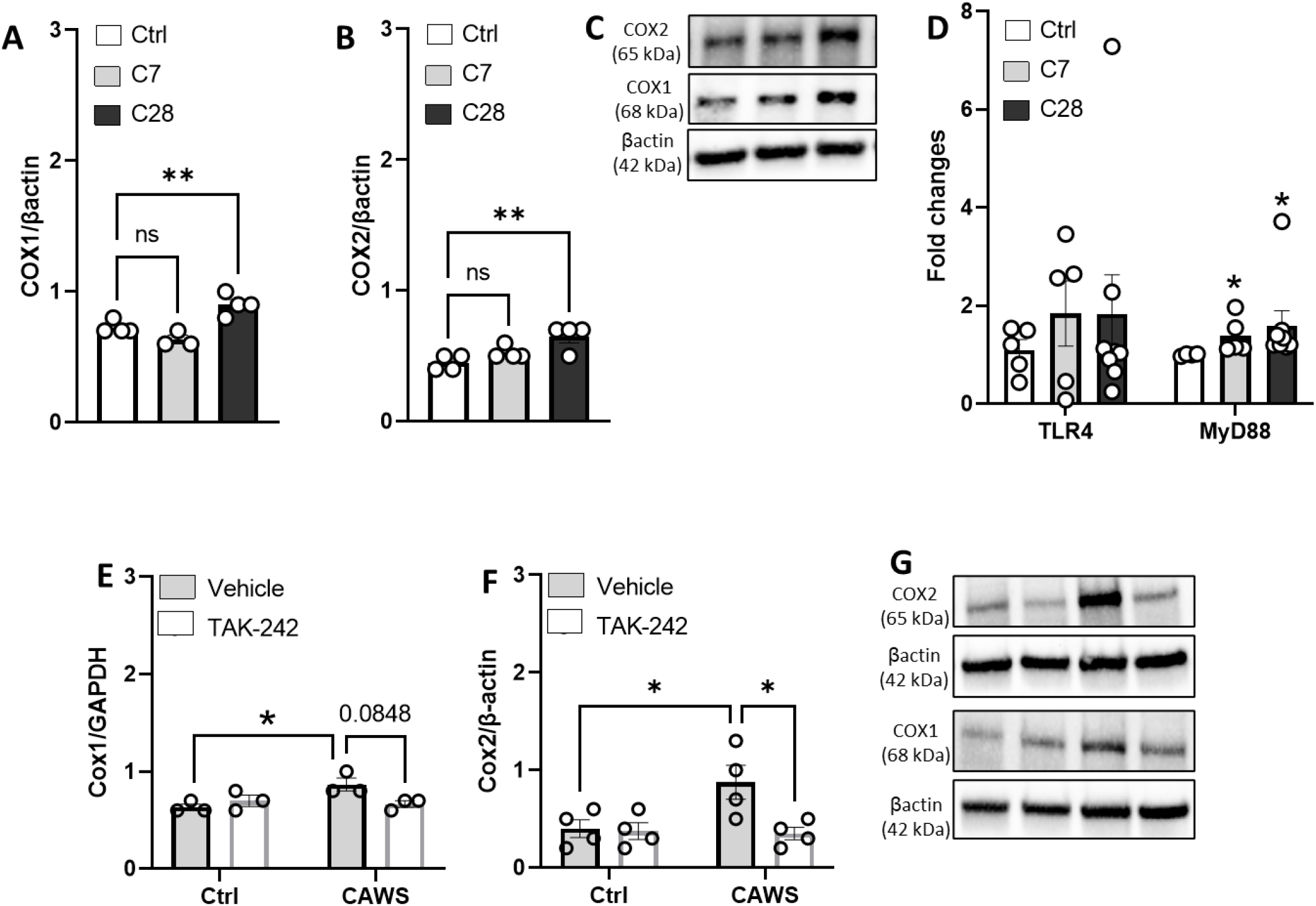
Vascular TLR4 may participate in CAWS-induced COXs activation. Effects of CAWS (10ug/mL, 0-24h) on COX1 (A and C) and COX2 (B and C) protein expression in Rat Aortic Smooth Muscle Cells (RASMC). Aortic gene expression of TLR4 and MyD88 (D) in samples from control (Ctrl): water injected for 5 days; CAWS 7 days (C7): CAWS injected for 5 days plus 2 additional days of wait; CAWS 28 days (C28): CAWS injected for 5 days plus 23 additional days of wait. RASMC treated with CAWS (10ug/mL) with or without TAK-242 (3_x_10^−5^M) on COX1 (E and G) and COX2 (F and G) protein expression. N=3-7. Data are presented as Mean ± SEM. *P<0.05 vs Ctrl in either, mouse samples or cells.

## Discussion

KD is the leading cause of acquired cardiovascular disease of unknown etiology that mostly affects infants and children under 5 years of age^3,4^ characterized by severe vasculitis, which can lead to coronary artery dilatations and aneurysms^1,3,4^. Thus, deciphering the cellular and molecular mechanisms by which arteries become injured is crucial. Herein, we characterized for the first time the vascular function and blood pressure regulation of a mouse model KD, as well as described COXs-derived products and ROS and NO bioavailability as possible dysfunctional mechanisms associated with vascular dysfunction. Finally, we demonstrated that the CAWS-induced KD model does not cause KDSS, at least not in our hands.

Patients with a history of KD present with vascular dysfunction^35,36^, which may persist over many years after resolution of KD^6,37^. Although the association between vascular dysfunction and KD has been almost unanimously reported before, few others have found that arteries from patients who recovered from KD are healthy and still working properly^5,8^. Furthermore, studies are mostly focused on evaluating the vascular function in patients who recovered from KD, rather than during the acute phase^6,8,34-36^. A limiting step for these studies is the lack of exploration on molecular mechanisms. Therefore, mouse models of KD are a useful approach to access the vascular function, as well as the possible mechanisms involved on vascular dysfunction associated with KD. Herein, we found that abdominal aortae from murine model of KD presents increased vascular contractility to noradrenaline in acute and convalescent phases.

COXs are key enzymes in arachidonic acid metabolism in cardiovascular physiology and pathophysiology^50,51^. Two major isoforms of COXs have been identified, COX1 and COX2. While COX1 is considered a constitutive isoform and responsible to produce its products within physiological environment, COX2 is thought to be an inducible isoform because of inflammatory process^52^ including atherosclerosis^53,54^ and hypertension^55^. COX1 also has been associated with cardiovascular diseases though. The relationship between COX2 and KD was previously observed in Chinese population by demonstrating that polymorphism on COX2 gene facilitates the KD appearance^56^. We observed that inhibition of COX1 and COX2, COX2 alone, or antagonism of TXA_2_R abolished the vascular hypercontractility in acute and convalescent phases of KD. TXA_2_ is a powerful agonist produced by COXs and a dysregulation of its levels or signaling has been described as a cardiovascular risk^50,51^. Yet, we confirmed that COX1 and 2 expressions, as well as TXA_2_R are elevated in abdominal aortae of our mouse model of KD. Aspirin has been used combined with IVIG treatment for KD because it can reduce pain and discomfort and help drop the fever^14^. The most recognized mechanism of action of aspirin is to block the synthesis of prostaglandins, for example TXA_2_, via inhibiting COXs^57^. Therefore, the use of aspirin in clinic might alleviate the vascular dysfunction in acute phase by blocking the COXs and TXA_2_ synthase. Interestingly, we also observed that VSMC per se can recognize the antigen (CAWS treatment) and translate COX1 and 2 expressions on a TLR4-dependent manner. TLR4 was discovered to be a crucial receptor in sterile inflammation via modulating COXs expression and TXA2, in addition to playing a highly specific role in immunological responses and KD ^58,59^,. Therefore, KD is associated with vascular dysfunction and upregulation of COXs, at least in part, via vascular TLR4.

An increase in oxidant status has been linked to numerous cardiovascular problems. ^30,31,60^, but whether changes in vascular redox signaling participates in vascular injury in KD is unknown. Infants with 2-10 years of age, diagnosed with medium-to-giant coronary aneurysms because of KD, showed improved endothelial function and decreased low-grade chronic inflammation after treatment with pravastatin, a 3-hydroxy-3-methylglutaryl-coenzyme (HMG-CoA) reductase inhibitor^34^. We and others have demonstrated that statins present pleiotropic effects (independent on lowering cholesterol levels) by acting as a potent antioxidant, blocking Noxs, and increasing NO production^30,61^. which results in cardiovascular protection including improvement on vascular function. We did not see difference in, but we found that Nox1 and 2 are elevated in abdominal aortae in C7 and C28. After pre-treating the aortae (during studies of vascular function) with an antioxidant agent (tempol), we abolished the vascular dysfunction in KD indicating that increase in ROS is leading to vascular dysfunction in KD. Interestingly, CAWS treatment only increased Nox1 in the C28 group’s aorta; it did not do so in isolated VSMC. Infiltrated immune cells, which are evidently more numerous during the convalescent phase, may be generating cytokines and raising Nox1. (Fig. 1). We recently discovered that RANTES (CCR5 ligand) leads to vascular inflammation and proliferation via Nox1. Additionally, it has previously been observed that TNFα^62^, IL6, and IL1β^63^ stimulate Nox1, which are elevated in the vasculature in our study.

Finally, we did not use any specific inhibitors for Nox1 or 2, but their upregulation may implicate that Noxs are producing ROS and contributing to vascular dysfunction in KD. Increase in ROS, specifically superoxide, may decrease NO bioavailability by forming peroxinitrite, thus reducing the vasorelaxant property of NO^64,65^. Our data indicate that there is no difference in NOS activity, but pharmacological inhibition of NOS with L-NAME blunted the difference in vascular constriction between control group and KD, which may suggest that NO bioavailability is reduced in abdominal aortae from KD.

KDSS refers to KD patients who have a greater than 20% drop in systolic blood pressure. Typically, these patients have elevated cytokine levels, are more likely to develop IVIG resistance, and have poorer cardiovascular outcomes than KD patients without KDSS^10,15,16^. We observed that CAWS murine model induced a mild, but not significant, decrease in blood pressure or heart rate, suggesting that, at least in our hand, this model is not appropriate to study KDSS. Perhaps if we have a larger cohort followed by a strong correlation between high cytokines levels (index of KDSS^10,15,16^) and blood pressure changes could be an alternative approach to further investigate the potential use of murine model of KD and KDSS.

In summary, we observed that KD, in acute and convalescent phases, is associated with vascular dysfunction characterized by hypercontractility to noradrenaline in abdominal aortae. Such change occurs dependent on increase in COX2-derived products and ROS and reduced NO bioavailability. Furthermore, KD inducer can increase COX2 expression directly in the vasculature by TLR4 signaling pathway (graphical abstract). Finally, refuting our initial hypothesis, the CAWS murine model does not induce significant changes in the blood pressure. We provide novel insights on the vascular mechanisms contributing to the vascular dysfunction in KD and place endothelial factors as attractive pharmacological targets to alleviate the vascular injury in KD. Future experiments are necessary to understand whether blockage of these factors (COXs and ROS) can confer better cardiovascular outcomes in children who had KD and whether vascular dysfunction is the trigger of vasculitis in KD.

## Authorship contribution statement

Mia DelVechio: Methodology, Formal analysis, Data curation, writing – original draft, writing – review & editing; Juliano V Alves: Methodology, Formal analysis, Data curation, Review & editing; Alina Z Saiyi: Methodology^;^ Rohit Goru: Methodology; Shubhnita Singh: Methodology, writing – review & editing; Joseph Galley: Methodology; Wanessa M Awata: Methodology, writing – review & editing; Ariane Bruder-Nascimento: Methodology, data curation, writing – original draft, writing – review & editing; Thiago Bruder-Nascimento: Conceptualization, Methodology, Validation, Formal analysis, Resources, Data curation, Writing – original draft, Writing – review & editing, Supervision, Project administration, Funding acquisition.

## Declaration of Competing Interest

The authors declare that they have no known competing financial interests or personal relationships that could have appeared to influence the work reported in this paper.

## Acknowledgments

This work was supported by: NHLBI-R00 (R00HL14013903), AHA-CDA (CDA857268), Vascular Medicine Institute, the Hemophilia Center of Western Pennsylvania Vitalant, Children’s Hospital of Pittsburgh of the UPMC Health System, and startup funds from University of Pittsburgh to TBN. All figures were designed using BioRender.

## Notes

### Competing Interest Statement

The authors have declared no competing interest.

## References

1 Noval Rivas, M. & Arditi, M. Kawasaki disease: pathophysiology and insights from mouse models. Nat Rev Rheumatol 16, 391–405, doi:10.1038/s41584-020-0426-0 (2020).

2 Naoe, S., Takahashi, K., Masuda, H. & Tanaka, N. Kawasaki disease. With particular emphasis on arterial lesions. Acta Pathol Jpn 41, 785–797, doi:10.1111/j.1440-1827.1991.tb01620.x (1991).

3 Sundel, R. P. Kawasaki disease. Rheum Dis Clin North Am 41, 63-73, viii, doi:10.1016/j.rdc.2014.09.010 (2015).

4 Uehara, R. & Belay, E. D. Epidemiology of Kawasaki disease in Asia, Europe, and the United States. J Epidemiol 22, 79–85, doi:10.2188/jea.je20110131 (2012).

5 Minich, L. L. et al. Delayed diagnosis of Kawasaki disease: what are the risk factors? Pediatrics 120, e1434–1440, doi:10.1542/peds.2007-0815 (2007).

6 Dhillon, R. et al. Endothelial dysfunction late after Kawasaki disease. Circulation 94, 2103–2106, doi:10.1161/01.cir.94.9.2103 (1996).

7 Fujiwara, H., Kawai, C. & Hamashima, Y. Clinicopathologic study of the conduction systems in 10 patients with Kawasaki’s disease (mucocutaneous lymph node syndrome). Am Heart J 96, 744–750, doi:10.1016/0002-8703(78)90007-8 (1978).

8 McCrindle, B. W., McIntyre, S., Kim, C., Lin, T. & Adeli, K. Are patients after Kawasaki disease at increased risk for accelerated atherosclerosis? J Pediatr 151, 244-248, 248 e241, doi:10.1016/j.jpeds.2007.03.056 (2007).

9 McCrindle, B. W. et al. Diagnosis, Treatment, and Long-Term Management of Kawasaki Disease: A Scientific Statement for Health Professionals From the American Heart Association. Circulation 135, e927–e999, doi:10.1161/CIR.0000000000000484 (2017).

10 Kanegaye, J. T. et al. Recognition of a Kawasaki disease shock syndrome. Pediatrics 123, e783–789, doi:10.1542/peds.2008-1871 (2009).

11 Liu, G., Wang, S. & Du, Z. Risk Factors of Intravenous Immunoglobulin Resistance in Children With Kawasaki Disease: A Meta-Analysis of Case-Control Studies. Front Pediatr 8, 187, doi:10.3389/fped.2020.00187 (2020).

12 Tremoulet, A. H. et al. Resistance to intravenous immunoglobulin in children with Kawasaki disease. J Pediatr 153, 117–121, doi:10.1016/j.jpeds.2007.12.021 (2008).

13 Turkucar, S. et al. Risk factors of intravenous immunoglobulin resistance and coronary arterial lesions in Turkish children with Kawasaki disease. Turk J Pediatr 62, 1–9, doi:10.24953/turkjped.2020.01.001 (2020).

14 Wallace, C. A., French, J. W., Kahn, S. J. & Sherry, D. D. Initial intravenous gammaglobulin treatment failure in Kawasaki disease. Pediatrics 105, E78, doi:10.1542/peds.105.6.e78 (2000).

15 Gamez-Gonzalez, L. B. et al. Kawasaki disease shock syndrome: Unique and severe subtype of Kawasaki disease. Pediatr Int 60, 781–790, doi:10.1111/ped.13614 (2018).

16 Li, Y. et al. Kawasaki disease shock syndrome: clinical characteristics and possible use of IL-6, IL-10 and IFN-gamma as biomarkers for early recognition. Pediatr Rheumatol Online J 17, 1, doi:10.1186/s12969-018-0303-4 (2019).

17 Anzai, F. et al. Crucial role of NLRP3 inflammasome in a murine model of Kawasaki disease. J Mol Cell Cardiol 138, 185–196, doi:10.1016/j.yjmcc.2019.11.158 (2020).

18 Hara, T. et al. Kawasaki disease: a matter of innate immunity. Clin Exp Immunol 186, 134–143, doi:10.1111/cei.12832 (2016).

19 Marek-Iannucci, S. et al. Autophagy-mitophagy induction attenuates cardiovascular inflammation in a murine model of Kawasaki disease vasculitis. JCI Insight 6, doi:10.1172/jci.insight.151981 (2021).

20 Miyabe, C. et al. Dectin-2-induced CCL2 production in tissue-resident macrophages ignites cardiac arteritis. J Clin Invest 129, 3610–3624, doi:10.1172/JCI123778 (2019).

21 Stock, A. T., Hansen, J. A., Sleeman, M. A., McKenzie, B. S. & Wicks, I. P. GM-CSF primes cardiac inflammation in a mouse model of Kawasaki disease. J Exp Med 213, 1983–1998, doi:10.1084/jem.20151853 (2016).

22 Tanaka, H. et al. Coronary Vasculitis Induced in Mice by Cell Wall Mannoprotein Fractions of Clinically Isolated Candida Species. Med Mycol J 61, 33–48, doi:10.3314/mmj.20-00008 (2020).

23 Bruder-Nascimento, T. et al. NLRP3 Inflammasome Mediates Aldosterone-Induced Vascular Damage. Circulation 134, 1866–1880, doi:10.1161/CIRCULATIONAHA.116.024369 (2016).

24 da Costa, R. M. et al. TNF-alpha induces vascular insulin resistance via positive modulation of PTEN and decreased Akt/eNOS/NO signaling in high fat diet-fed mice. Cardiovasc Diabetol 15, 119, doi:10.1186/s12933-016-0443-0 (2016).

25 Itani, H. A. et al. Activation of Human T Cells in Hypertension: Studies of Humanized Mice and Hypertensive Humans. Hypertension 68, 123–132, doi:10.1161/HYPERTENSIONAHA.116.07237 (2016).

26 Cau, S. B. et al. Angiotensin-II activates vascular inflammasome and induces vascular damage. Vascul Pharmacol 139, 106881, doi:10.1016/j.vph.2021.106881 (2021).

27 Nedeljkovic, Z. S., Gokce, N. & Loscalzo, J. Mechanisms of oxidative stress and vascular dysfunction. Postgrad Med J 79, 195-199; quiz 198-200, doi:10.1136/pmj.79.930.195 (2003).

28 Davignon, J. & Ganz, P. Role of endothelial dysfunction in atherosclerosis. Circulation 109, III27–32, doi:10.1161/01.CIR.0000131515.03336.f8 (2004).

29 Brandes, R. P. Endothelial dysfunction and hypertension. Hypertension 64, 924–928, doi:10.1161/HYPERTENSIONAHA.114.03575 (2014).

30 Bruder-Nascimento, T. et al. Atorvastatin inhibits pro-inflammatory actions of aldosterone in vascular smooth muscle cells by reducing oxidative stress. Life Sci 221, 29–34, doi:10.1016/j.lfs.2019.01.043 (2019).

31 Bruder-Nascimento, T. et al. Angiotensin II induces Fat1 expression/activation and vascular smooth muscle cell migration via Nox1-dependent reactive oxygen species generation. J Mol Cell Cardiol 66, 18–26, doi:10.1016/j.yjmcc.2013.10.013 (2014).

32 Bruder-Nascimento, T. et al. Leptin Restores Endothelial Function via Endothelial PPARgamma-Nox1-Mediated Mechanisms in a Mouse Model of Congenital Generalized Lipodystrophy. Hypertension 74, 1399–1408, doi:10.1161/HYPERTENSIONAHA.119.13398 (2019).

33 Singh, S., Bruder-Nascimento, A., Belin de Chantemele, E. J. & Bruder-Nascimento, T. CCR5 antagonist treatment inhibits vascular injury by regulating NADPH oxidase 1. Biochem Pharmacol 195, 114859, doi:10.1016/j.bcp.2021.114859 (2022).

34 Duan, C., Du, Z. D., Wang, Y. & Jia, L. Q. Effect of pravastatin on endothelial dysfunction in children with medium to giant coronary aneurysms due to Kawasaki disease. World J Pediatr 10, 232–237, doi:10.1007/s12519-014-0498-5 (2014).

35 Niboshi, A., Hamaoka, K., Sakata, K. & Yamaguchi, N. Endothelial dysfunction in adult patients with a history of Kawasaki disease. Eur J Pediatr 167, 189–196, doi:10.1007/s00431-007-0452-9 (2008).

36 Borzutzky, A. et al. High sensitivity C-reactive protein and endothelial function in Chilean patients with history of Kawasaki disease. Clin Rheumatol 27, 845–850, doi:10.1007/s10067-007-0808-6 (2008).

37 Ikemoto, Y., Ogino, H., Teraguchi, M. & Kobayashi, Y. Evaluation of preclinical atherosclerosis by flow-mediated dilatation of the brachial artery and carotid artery analysis in patients with a history of Kawasaki disease. Pediatr Cardiol 26, 782–786, doi:10.1007/s00246-005-0921-8 (2005).

38 Chen, S. et al. Marked acceleration of atherosclerosis after Lactobacillus casei-induced coronary arteritis in a mouse model of Kawasaki disease. Arterioscler Thromb Vasc Biol 32, e60–71, doi:10.1161/ATVBAHA.112.249417 (2012).

39 Silva, A. A. et al. Cardiovascular risk factors after Kawasaki disease: a case-control study. J Pediatr 138, 400–405, doi:10.1067/mpd.2001.111430 (2001).

40 Nagi-Miura, N. et al. Lethal and severe coronary arteritis in DBA/2 mice induced by fungal pathogen, CAWS, Candida albicans water-soluble fraction. Atherosclerosis 186, 310–320, doi:10.1016/j.atherosclerosis.2005.08.014 (2006).

41 Tada, R., Nagi-Miura, N., Adachi, Y. & Ohno, N. The influence of culture conditions on vasculitis and anaphylactoid shock induced by fungal pathogen Candida albicans cell wall extract in mice. Microb Pathog 44, 379–388, doi:10.1016/j.micpath.2007.10.013 (2008).

42 Shepherd, M. G. & Sullivan, P. A. The production and growth characteristics of yeast and mycelial forms of Candida albicans in continuous culture. J Gen Microbiol 93, 361–370, doi:10.1099/00221287-93-2-361 (1976).

43 Bruder-Nascimento, T. et al. Deletion of protein tyrosine phosphatase 1b in proopiomelanocortin neurons reduces neurogenic control of blood pressure and protects mice from leptin- and sympatho-mediated hypertension. Pharmacol Res 102, 235–244, doi:10.1016/j.phrs.2015.10.012 (2015).

44 Bruder-Nascimento, T., Ekeledo, O. J., Anderson, R., Le, H. B. & Belin de Chantemele, E. J. Long Term High Fat Diet Treatment: An Appropriate Approach to Study the Sex-Specificity of the Autonomic and Cardiovascular Responses to Obesity in Mice. Front Physiol 8, 32, doi:10.3389/fphys.2017.00032 (2017).

45 Zheng, Y. et al. Melatonin alleviates vascular endothelial cell damage by regulating an autophagy-apoptosis axis in Kawasaki disease. Cell Prolif 55, e13251, doi:10.1111/cpr.13251 (2022).

46 Bruder-Nascimento, T. et al. Ptp1b deletion in pro-opiomelanocortin neurons increases energy expenditure and impairs endothelial function via TNF-alpha dependent mechanisms. Clin Sci (Lond) 130, 881–893, doi:10.1042/CS20160073 (2016).

47 Kawasaki, A. & Needleman, P. Contribution of thromboxane to renal resistance changes in the isolated perfused hydronephrotic rabbit kidney. Circ Res 50, 486–490, doi:10.1161/01.res.50.4.486 (1982).

48 Hernanz, R. et al. Toll-like receptor 4 contributes to vascular remodelling and endothelial dysfunction in angiotensin II-induced hypertension. Br J Pharmacol 172, 3159–3176, doi:10.1111/bph.13117 (2015).

49 De Batista, P. R. et al. Toll-like receptor 4 upregulation by angiotensin II contributes to hypertension and vascular dysfunction through reactive oxygen species production. PLoS One 9, e104020, doi:10.1371/journal.pone.0104020 (2014).

50 Schjerning, A. M., McGettigan, P. & Gislason, G. Cardiovascular effects and safety of (non-aspirin) NSAIDs. Nat Rev Cardiol 17, 574–584, doi:10.1038/s41569-020-0366-z (2020).

51 Antman, E. M., DeMets, D. & Loscalzo, J. Cyclooxygenase inhibition and cardiovascular risk. Circulation 112, 759–770, doi:10.1161/CIRCULATIONAHA.105.568451 (2005).

52 Kirkby, N. S. et al. Cyclooxygenase-1, not cyclooxygenase-2, is responsible for physiological production of prostacyclin in the cardiovascular system. Proc Natl Acad Sci U S A 109, 17597–17602, doi:10.1073/pnas.1209192109 (2012).

53 Pratico, D., Tillmann, C., Zhang, Z. B., Li, H. & FitzGerald, G. A. Acceleration of atherogenesis by COX-1-dependent prostanoid formation in low density lipoprotein receptor knockout mice. Proc Natl Acad Sci U S A 98, 3358–3363, doi:10.1073/pnas.061607398 (2001).

54 Burleigh, M. E. et al. Cyclooxygenase-2 promotes early atherosclerotic lesion formation in LDL receptor-deficient mice. Circulation 105, 1816–1823, doi:10.1161/01.cir.0000014927.74465.7f (2002).

55 Bomfim, G. F. et al. Toll-like receptor 4 contributes to blood pressure regulation and vascular contraction in spontaneously hypertensive rats. Clin Sci (Lond) 122, 535–543, doi:10.1042/CS20110523 (2012).

56 Li, S. et al. The Relationship of COX-2 Gene Polymorphisms and Susceptibility to Kawasaki Disease in Chinese Population. Immunol Invest 48, 181–189, doi:10.1080/08820139.2018.1529790 (2019).

57 Flower, R. What are all the things that aspirin does? BMJ 327, 572–573, doi:10.1136/bmj.327.7415.572 (2003).

58 Firmal, P., Shah, V. K. & Chattopadhyay, S. Insight Into TLR4-Mediated Immunomodulation in Normal Pregnancy and Related Disorders. Front Immunol 11, 807, doi:10.3389/fimmu.2020.00807 (2020).

59 Nakashima, T. et al. TLR4 is a critical regulator of angiotensin II-induced vascular remodeling: the roles of extracellular SOD and NADPH oxidase. Hypertens Res 38, 649–655, doi:10.1038/hr.2015.55 (2015).

60 Harvey, A., Montezano, A. C., Lopes, R. A., Rios, F. & Touyz, R. M. Vascular Fibrosis in Aging and Hypertension: Molecular Mechanisms and Clinical Implications. Can J Cardiol 32, 659–668, doi:10.1016/j.cjca.2016.02.070 (2016).

61 Bruder-Nascimento, T. et al. Vascular injury in diabetic db/db mice is ameliorated by atorvastatin: role of Rac1/2-sensitive Nox-dependent pathways. Clin Sci (Lond) 128, 411–423, doi:10.1042/CS20140456 (2015).

62 de Jesus, D. S. et al. Nox1/Ref-1-mediated activation of CREB promotes Gremlin1-driven endothelial cell proliferation and migration. Redox Biol 22, 101138, doi:10.1016/j.redox.2019.101138 (2019).

63 Chamberlain, J. et al. Interleukin-1 regulates multiple atherogenic mechanisms in response to fat feeding. PLoS One 4, e5073, doi:10.1371/journal.pone.0005073 (2009).

64 Loperena, R. & Harrison, D. G. Oxidative Stress and Hypertensive Diseases. Med Clin North Am 101, 169–193, doi:10.1016/j.mcna.2016.08.004 (2017).

65 Dikalova, A. E. et al. Therapeutic targeting of mitochondrial superoxide in hypertension. Circ Res 107, 106–116, doi:10.1161/CIRCRESAHA.109.214601 (2010).

